# Midkine and the immunosuppressive tumor microenvironment in pancreatic ductal adenocarcinoma: evidence from Mendelian randomization, multi-omics factor analysis, and single-cell transcriptomics

**DOI:** 10.64898/2026.11.12.766325

**Authors:** Zhengchen Yang, Xuejian Chen, Zihao Hu, Lishan Xu

## Abstract

Pancreatic ductal adenocarcinoma (PDAC) remains one of the most lethal malignancies, in part because its tumor microenvironment actively suppresses antitumor immunity. Midkine (MDK) is a secreted heparin-binding growth factor that is overexpressed in a majority of PDAC tumors, but whether it contributes causally to disease risk and through which cell types it acts have not been firmly established. We combined cis-eQTL Mendelian randomization (GTEx pancreas, n = 328 donors; FinnGen PDAC, 1,252 cases) with multi-omics factor analysis (MOFA) of TCGA-PAAD tumors (n = 149), single-sample gene set enrichment analysis (ssGSEA), cross-species single-cell RNA sequencing, and DepMap CRISPR dependency screening. Mendelian randomization identified MDK as the only gene passing false discovery rate correction for PDAC risk (inverse-variance-weighted odds ratio 1.51, 95% CI 1.19–1.91, p = 6.7 × 10⁻⁴; FDR = 0.008), with concordant direction across five methods and five independent GWAS. MDK was overexpressed in tumors relative to adjacent normal tissue in an independent cohort (GSE62452, n = 130, p = 5 × 10⁻⁸), and high expression was associated with shorter overall survival (10.2 versus 21.1 months, p = 0.023). In the MOFA decomposition, MDK localized to a factor characterized by tertiary lymphoid structure gene expression but negative correlation with dendritic cells and macrophages, a pattern consistent with immune exclusion. Single-cell analysis and published functional studies indicated that MDK is produced by cancer-associated fibroblasts and endothelial cells rather than by tumor epithelium, and DepMap data confirmed that MDK is not a cell-autonomous fitness gene. Collectively, these results support a model in which MDK acts through the stromal-immune compartment rather than through tumor cell-intrinsic mechanisms, and raise the possibility that MDK blockade could restore antitumor immunity in PDAC.

## Introduction

Pancreatic ductal adenocarcinoma (PDAC) has a five-year survival below 12% and is expected to become the second leading cause of cancer death by 2030 [1]. A defining feature of PDAC is its immunosuppressive tumor microenvironment, in which cytotoxic T cells are excluded and myeloid-derived suppressor cells and tumor-associated macrophages predominate [2,3,4]. This immune-excluded state is widely regarded as a principal reason that PDAC responds poorly to checkpoint blockade and other immunotherapies [9]. Identifying the molecules that maintain this state is therefore of practical importance.

Midkine (MDK) is a 13-kDa heparin-binding growth factor first isolated from retinoic acid-treated embryonal carcinoma cells [5]. It signals through syndecans, the receptor-type tyrosine phosphatase PTPRZ1, and integrins α4β1/α6β1, and it has been reported to promote proliferation, migration, angiogenesis, and cell survival [6]. Immunohistochemical studies have detected MDK protein in approximately 62% of PDAC tumors but in only 26% of adjacent normal pancreas [7], and MDK expression correlates with perineural invasion, advanced tumor stage, and reduced survival [8]. A substantial body of work in other tumor types has shown that MDK can impair dendritic cell function and polarize macrophages toward an immunosuppressive M2 phenotype [6,18].

Two questions remain unresolved. First, whether MDK upregulation is causally related to PDAC risk, or is instead a marker of aggressive disease. Second, which cell types produce MDK within the PDAC microenvironment and whether its effects are tumor cell-intrinsic or microenvironmental. In this study we address both questions by combining Mendelian randomization, multi-omics factor analysis, immune deconvolution, single-cell transcriptomics, and functional genomics. Mendelian randomization estimates causal effects under specific assumptions about the validity of the genetic instruments, and is used here to test whether MDK expression causally influences PDAC risk.

## Materials and Methods

### Cis-eQTL Mendelian randomization

Pancreatic cis-eQTL data were obtained from GTEx Analysis v8 (328 donors). For each of 34 pancreatic neuro-tumor interaction genes, the most significant cis-eQTL single nucleotide polymorphism (SNP) within ±1 Mb of the transcription start site (F-statistic > 10, p < 5 × 10⁻⁷) was selected as the instrumental variable. Effect alleles were assigned as the alternative allele, obtained from the fourth field of the GTEx variant_id. PDAC summary statistics were obtained from FinnGen Release 13 (ADENO_DUCTAL phenotype; 1,252 cases, 372,159 controls). Mendelian randomization relies on three assumptions for a valid instrumental variable: relevance, that the genetic variant is robustly associated with the exposure; independence, that the variant is not associated with confounders of the exposure–outcome relationship; and the exclusion restriction, that the variant affects the outcome only through the exposure. Variants with F-statistic > 10 were selected to satisfy the relevance assumption. Because the independence and exclusion-restriction assumptions cannot be fully verified empirically, we applied methods with differing sensitivity to horizontal pleiotropy (weighted median, MR-Egger, and single-SNP Wald) together with the Steiger directionality test to assess their plausibility. As the exposure and outcome were drawn from independent samples (GTEx and FinnGen), there was no sample overlap. Five methods were applied following current guidelines [22]: inverse-variance-weighted (IVW) as the primary analysis, together with weighted median, simple mode, MR-Egger, and single-SNP Wald ratio. Benjamini-Hochberg correction was applied across 12 IVW tests. Heterogeneity among instrument-specific estimates was assessed with Cochran’s Q statistic and the I² index, and leave-one-out analysis was performed by sequentially omitting each variant and recomputing the inverse-variance-weighted estimate. A post hoc power calculation for the primary MDK analysis was performed using the two-sample Mendelian randomization power formula of Burgess [25]; given the observed instrument strength (R² ≈ 8.5% for the three MDK instruments) and 1,252 PDAC cases, the minimum odds ratio detectable with 80% power was estimated. Colocalization was assessed with the coloc package (v5.2.3) using default priors. Replication was attempted in four additional PDAC GWAS: FinnGen WIDE, PanScan I, BioBank Japan, and a 2021 East Asian GWAS. Analyses were performed in R 4.6.1 using the TwoSampleMR (0.7.9) package.

### TCGA-PAAD multi-omics data and MOFA

TCGA-PAAD data were downloaded from the cBioPortal Firehose Legacy dataset. RNA-seq RSEM values were log2(RSEM+1)-transformed; HM450 beta-values were averaged per gene; copy number was used as GISTIC2 discrete values; and non-synonymous mutations were binarized. The top 5,000 features by variance were retained per layer, and samples present in all four layers with survival annotation were retained (n = 149). Multi-omics factor analysis was performed with MOFA2 v1.22.0 (mofapy2 v0.7.4 backend) using 15 latent factors, Gaussian likelihoods for expression, methylation, and copy number, a Bernoulli likelihood for mutations, and 2,000 iterations. Convergence was verified by monitoring the evidence lower bound.

### Survival analysis

Univariate Cox proportional hazards regression was performed for each of the 15 MOFA factor scores using the survival package (v3.7). In addition, a DeepSurv network was implemented as a multilayer perceptron (15-16-8-1, ReLU activations, dropout 0.3, Adam optimizer) with a Cox partial likelihood loss, and was evaluated by 5-fold cross-validation using Harrell’s C-index. Random forest and elastic net Cox models were evaluated in the same fold structure as baselines.

### Immune deconvolution

Single-sample gene set enrichment analysis was performed with the GSVA package (v2.6.3) using the Bindea 2013 immune signature panel (27 cell types). Spearman correlations were computed between MDK expression and each immune cell score, and Wilcoxon tests compared MDK-high and MDK-low groups defined by median split.

### Single-cell RNA-seq

Mouse KPC PDAC single-cell data (3 PBS-treated samples, 9,000 cells) were obtained from Dryad (doi:10.5061/dryad.kkwh70shs). Human PDAC CD45+ immune cell data (5,000 cells) were obtained from GEO (GSE235449). Cells were classified using canonical markers, and MDK was quantified as log2(CPM+1). The proportion of MDK-positive cells was computed per cell type.

### DepMap CRISPR dependency and IHC

MDK Chronos gene effect scores were obtained from DepMap Public 25Q3 (1,208 cell lines). Pancreatic cancer lines (OncotreeCode PAAD/PAASC, n = 48) were compared with the remaining lines using Welch’s t-test, Cohen’s d, and the Mann-Whitney U test. MDK immunohistochemistry images (antibody CAB010055, 12 cases) were obtained from the Human Protein Atlas.

### Statistics

Analyses were performed in Python 3.13.12 (NumPy 2.5.1, SciPy 1.17.0, Pandas 3.0.5) and R 4.6.1. All tests were two-sided, and p < 0.05 was considered significant.

## Results

### Mendelian randomization identifies MDK as a causal PDAC risk gene

Mendelian randomization was performed for 34 pancreatic neuro-tumor interaction genes. MDK was the only gene to pass false discovery rate correction, with an inverse-variance-weighted odds ratio of 1.51 (95% CI 1.19–1.91, p = 6.7 × 10⁻⁴, FDR = 0.008; Fig 1A, S1 Table). Weighted median, simple mode, MR-Egger, and single-SNP Wald estimates were all directionally consistent, and Steiger filtering supported a causal direction from expression to disease (p = 8.3 × 10⁻⁹). In four additional PDAC GWAS (FinnGen WIDE, OR 1.22, p = 0.008; PanScan I; BioBank Japan; and a 2021 East Asian GWAS), no dataset indicated a protective direction for MDK. Colocalization gave a modest PP.H4 of 0.24, with a PP.H4/PP.H3 ratio of 6.8 favoring a shared causal variant; the low absolute value reflects limited GWAS power rather than evidence against colocalization [21]. Cochran’s Q indicated no significant heterogeneity among the three instruments (Q = 0.65, df = 2, p = 0.72; I² = 0%), and the MR-Egger intercept provided no evidence of directional horizontal pleiotropy (intercept −0.14, p = 0.57). Leave-one-out analysis showed that sequential omission of any single variant left the inverse-variance-weighted estimate directionally unchanged and nominally significant (p = 0.0011–0.0126).

**Fig 1.**
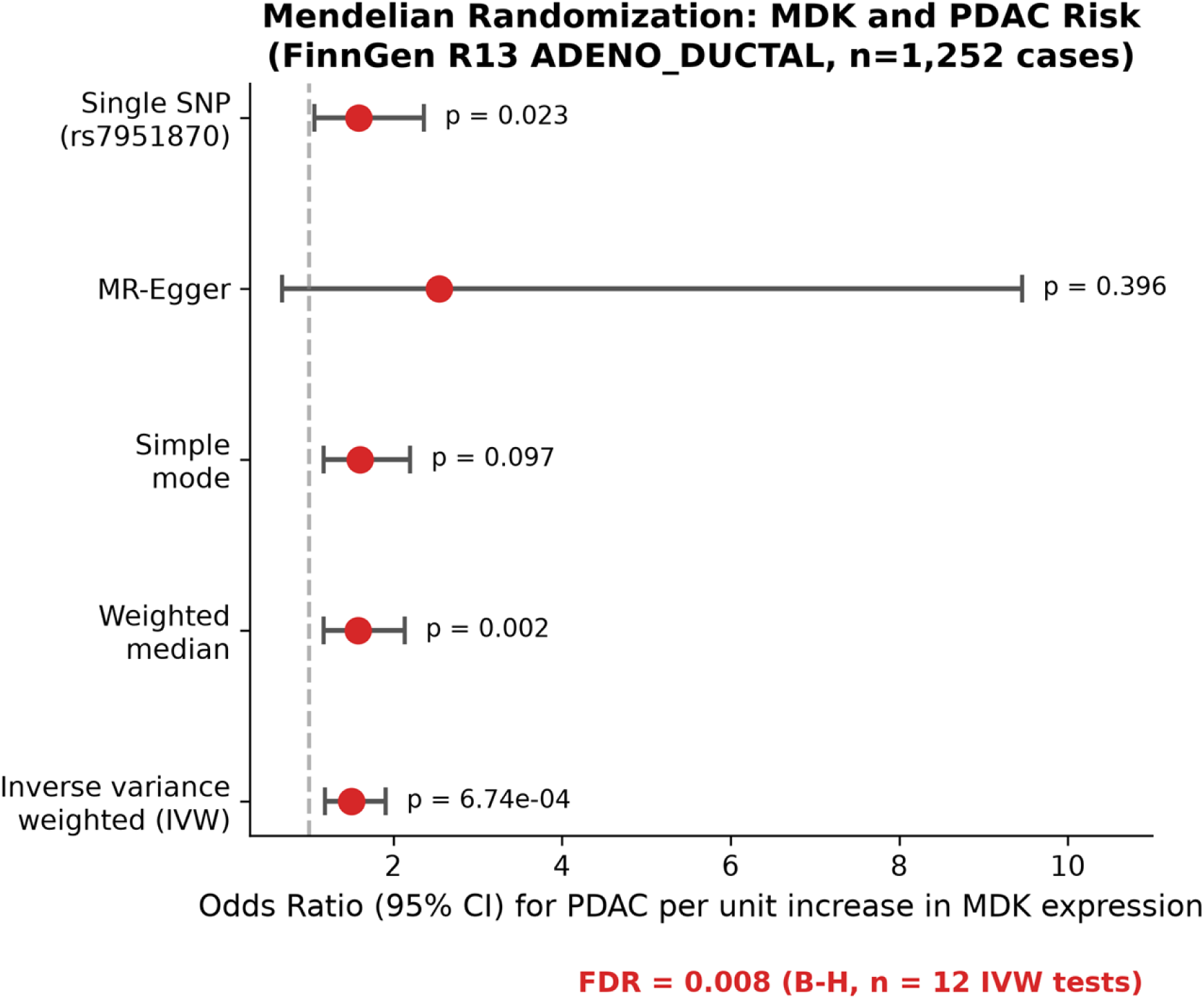
Mendelian randomization of MDK and pancreatic cancer risk. (A) Forest plot of odds ratios across five MR methods.

**Fig 1.**
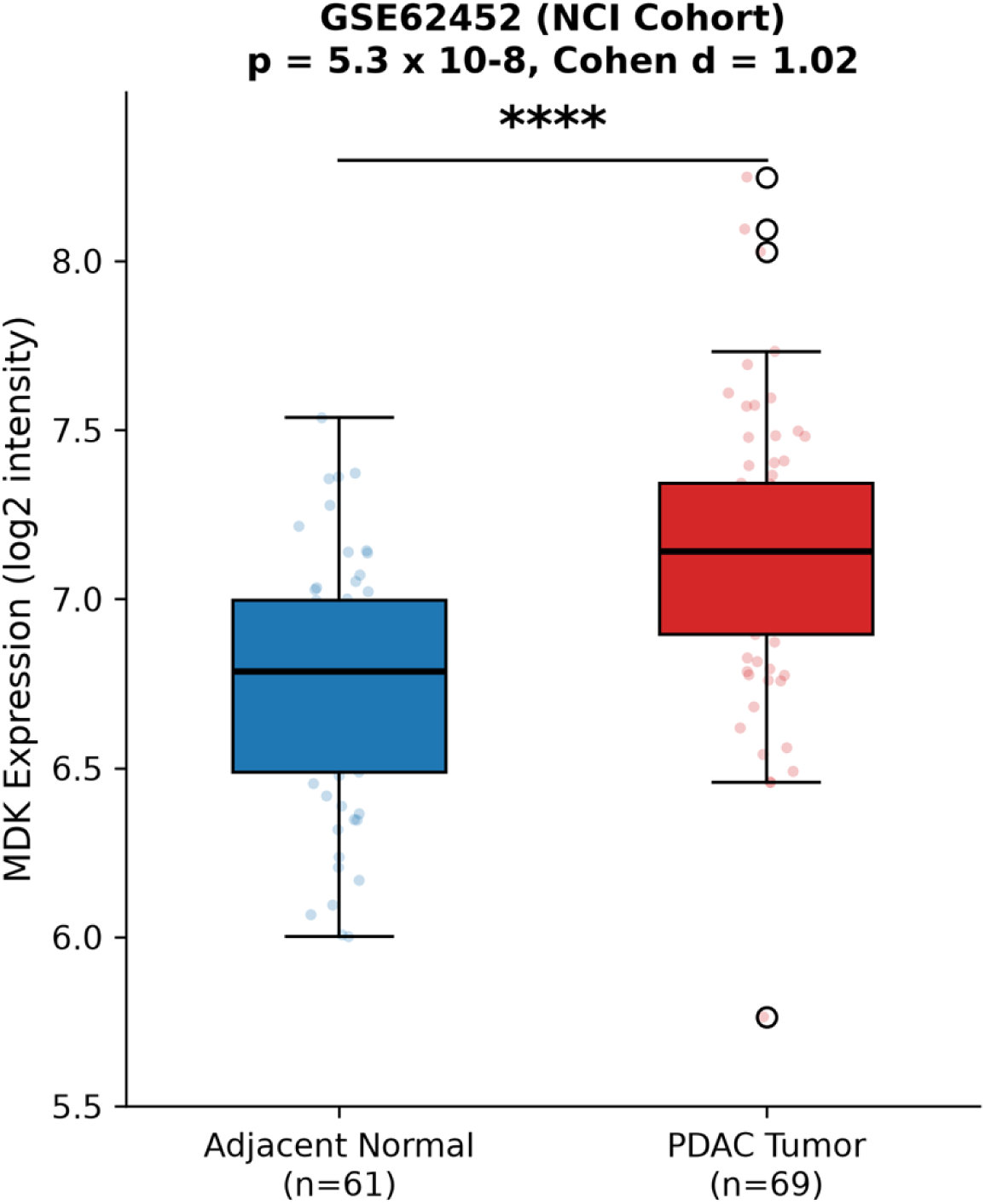
(B) MDK expression in tumor versus adjacent normal tissue in GSE62452.

**Fig 1.**
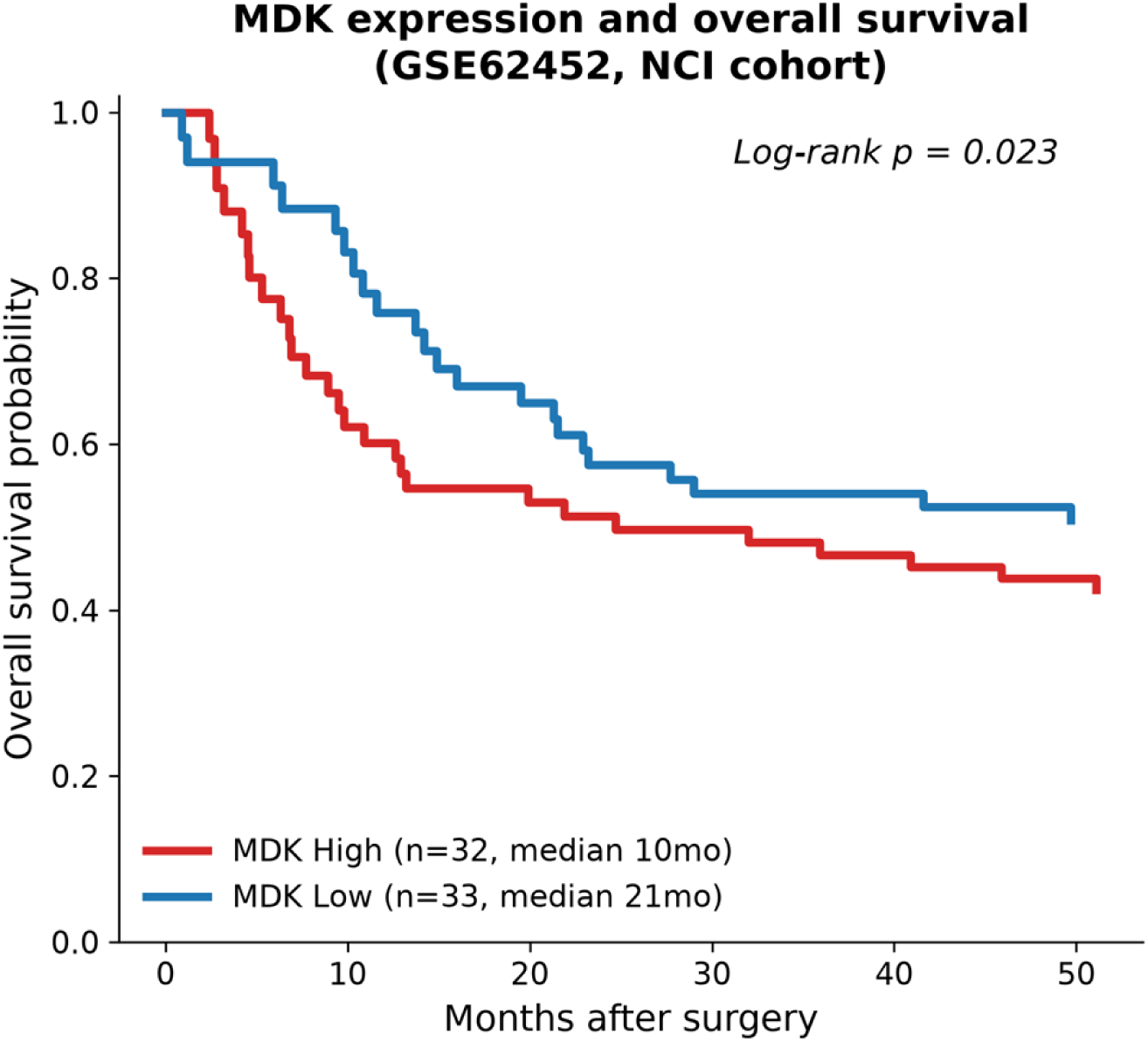
(C) Overall survival stratified by MDK expression in GSE62452.

We note one methodological point relevant to replication. The GTEx effect allele at MDK must be assigned as the alternative allele. Use of the reference allele inverted the estimated direction for all 29 genes analyzed; assignment of the alternative allele, validated against the independent GWAS, restored concordance.

### MDK is overexpressed in PDAC and associated with shorter survival

In an independent NCI cohort (GSE62452, 69 tumors and 61 adjacent normal samples), MDK expression was higher in tumors (mean 7.13 versus 6.74 log2 intensity, p = 5 × 10⁻⁸, Cohen’s d = 1.02; Fig 1B). At the protein level, CPTAC mass spectrometry confirmed MDK upregulation in PDAC (n = 235, p < 4 × 10⁻²¹), and Human Protein Atlas IHC showed positivity in 11 of 12 cases (Fig 5).

MDK expression was not associated with tumor stage (p = 0.80) or grade (p = 0.19; S1 Fig). However, high expression, defined by median split, was associated with shorter overall survival (10.2 versus 21.1 months, log-rank p = 0.023; Fig 1C). The corresponding analysis in TCGA-PAAD (n = 149) showed a concordant but non-significant trend (HR 1.06), and a fixed-effect meta-analysis of the two cohorts indicated no heterogeneity (I² = 0%), suggesting that the difference in significance reflects statistical precision rather than a biological inconsistency.

### MDK localizes to a TLS-enriched, immune-excluded MOFA factor

MOFA was applied to four omics layers of TCGA-PAAD. The 15-factor model converged, with copy number (49.9%) and methylation (37.2%) accounting for most of the explained variance (Fig 2A). MDK methylation contributed to Factor1 (weight−0.074) and MDK copy number to Factor5 (weight −0.115). MDK mRNA was not among the top 5,000 variable genes, indicating that its dysregulation is carried primarily by methylation and copy number.

**Fig 2.**
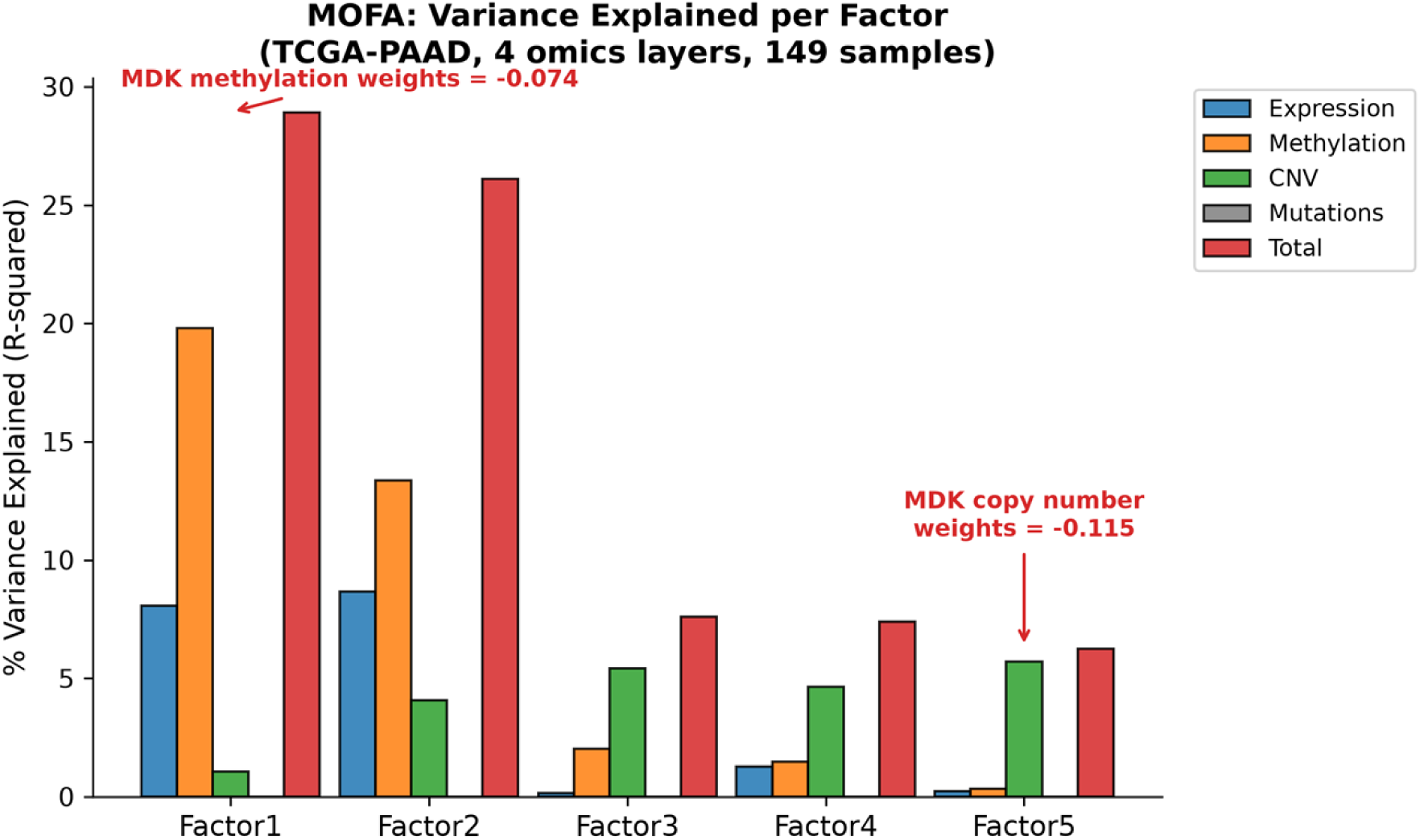
Multi-omics factor analysis of TCGA-PAAD. (A) Variance explained per factor across four omics layers.

**Fig 2.**
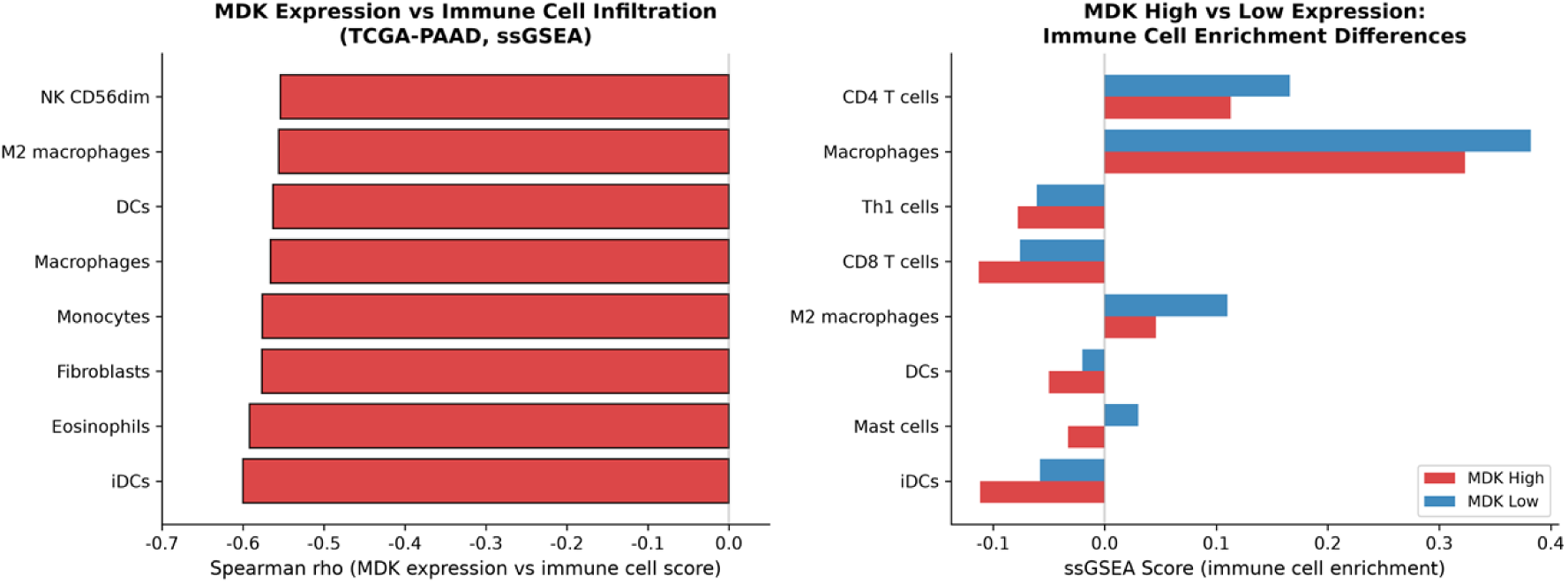
(B) Correlation between MDK expression and immune cell enrichment, and differences between MDK-high and MDK-low tumors.

Factor1 (R² = 28.9%) was characterized by expression of tertiary lymphoid structure (TLS) markers including CXCL13, TCL1A, and REG4, together with methylation features enriched in immune and mesenchymal pathways. At the same time, Factor1 was negatively correlated with dendritic cells (ρ = −0.60), macrophages (ρ = −0.57), and CD8+ T cells (ρ = −0.40) in immune deconvolution. This combination, TLS gene expression in the absence of antigen-presenting cell infiltration, matches the immune-excluded phenotype described in PDAC [3,13]. The presence of MDK within this factor is consistent with its reported ability to inhibit dendritic cell maturation and promote M2 polarization [6,18].

### MDK-high tumors are depleted of antigen-presenting cells

ssGSEA was performed with a 27-cell-type immune signature panel. Twenty of 27 cell types differed between MDK-high and MDK-low tumors (Wilcoxon p < 0.05; Fig 2B), with the largest reductions observed for immature dendritic cells, mast cells, conventional dendritic cells, and M2 macrophages. MDK was inversely correlated with SMAD4 at the transcript level (ρ = −0.42), and tumors with biallelic SMAD4 inactivation (mutation or deep deletion, n = 59) had higher MDK than SMAD4-intact tumors (p = 0.022, Cohen’s d = 0.37; S2 Fig). This links MDK upregulation to inactivation of the TGF-β pathway, an event with established immunosuppressive consequences in PDAC [9].

### Single-cell analysis localizes MDK to stromal and myeloid compartments

In mouse KPC tumors, MDK detection was sparse (0.6% of cells), as expected for a secreted factor with high dropout in droplet-based single-cell RNA-seq. Among positive cells, MDK was more frequent in cancer-associated fibroblasts (1.4%) and endothelial cells (1.5%) than in tumor epithelial cells (0.3%; Fig 3A). In human PDAC CD45+ immune cells, MDK positivity was highest in monocytes (7.8%) and macrophages (5.9%), with little expression in T and NK cells (Fig 3B).

**Fig 3.**
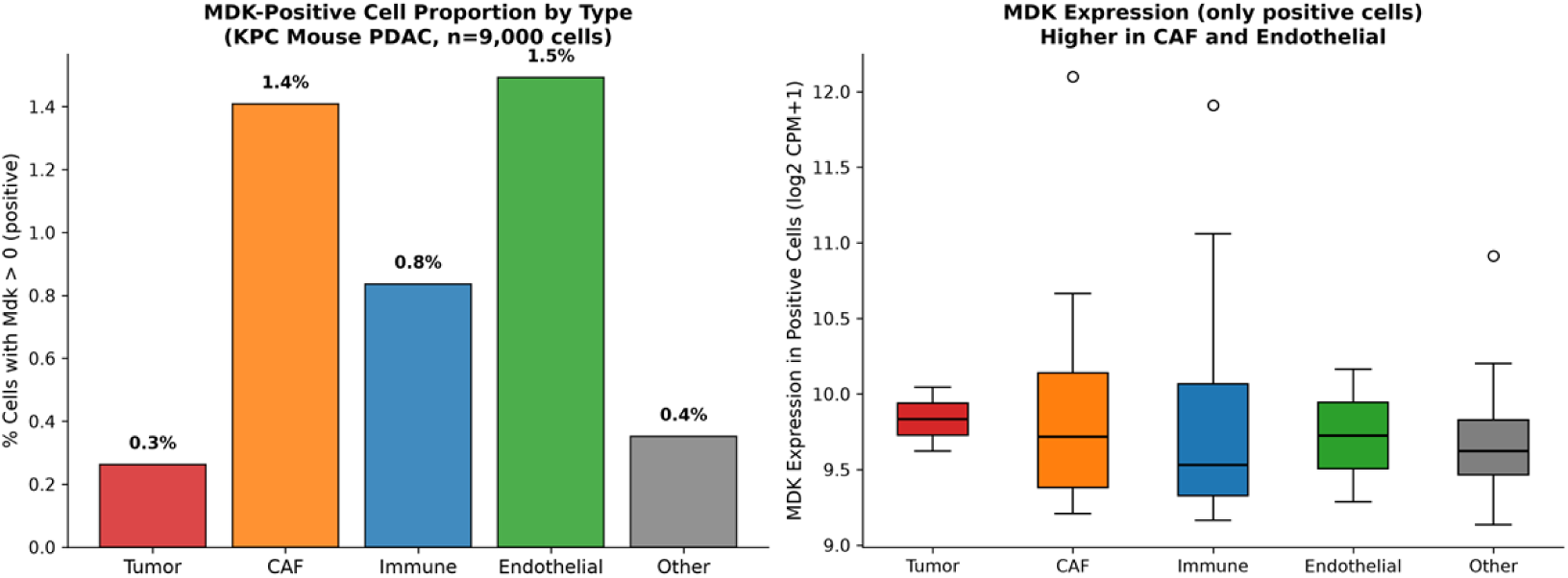
Single-cell RNA-seq of MDK. (A) Mouse KPC tumors; (B) human PDAC CD45+ immune cells.

Because single-cell dropout limits inference for secreted proteins, we also considered published functional evidence. MDK secretion by patient-derived cancer-associated fibroblasts has been demonstrated directly [15]; a PDAC-targeted delivery platform has exploited stromal MDK overexpression as a homing ligand [16]; and in esophageal carcinoma, a specific fibroblast subset has been shown to signal to tumor cells through an MDK–SDC1 axis [17]. Together with the absence of a tumor cell fitness effect in DepMap, these observations point to stromal cells, in particular cancer-associated fibroblasts, as the principal source of MDK.

### MDK is not a cell-autonomous fitness gene

In DepMap (1,208 cell lines), no line showed MDK essentiality (Chronos < −1.0) and only 1.4% reached moderate dependency (Chronos < −0.5; Fig 4). Pancreatic cancer lines had weaker MDK dependency than other lineages (mean −0.063 versus −0.104, p = 0.037, Cohen’s d = 0.28). The absence of a fitness effect, combined with the causal expression-to-disease association, is consistent with a microenvironmental rather than cell-autonomous mode of action.

**Fig 4.**
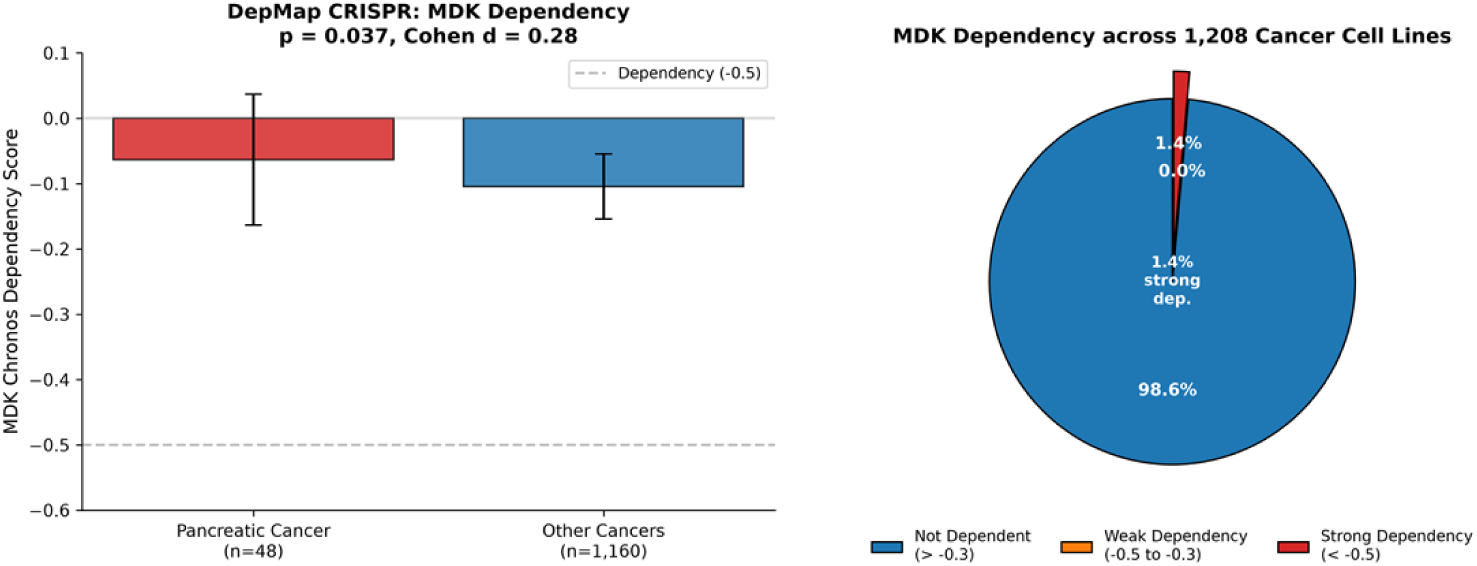
DepMap CRISPR dependency scores for MDK.

**Fig 5.**
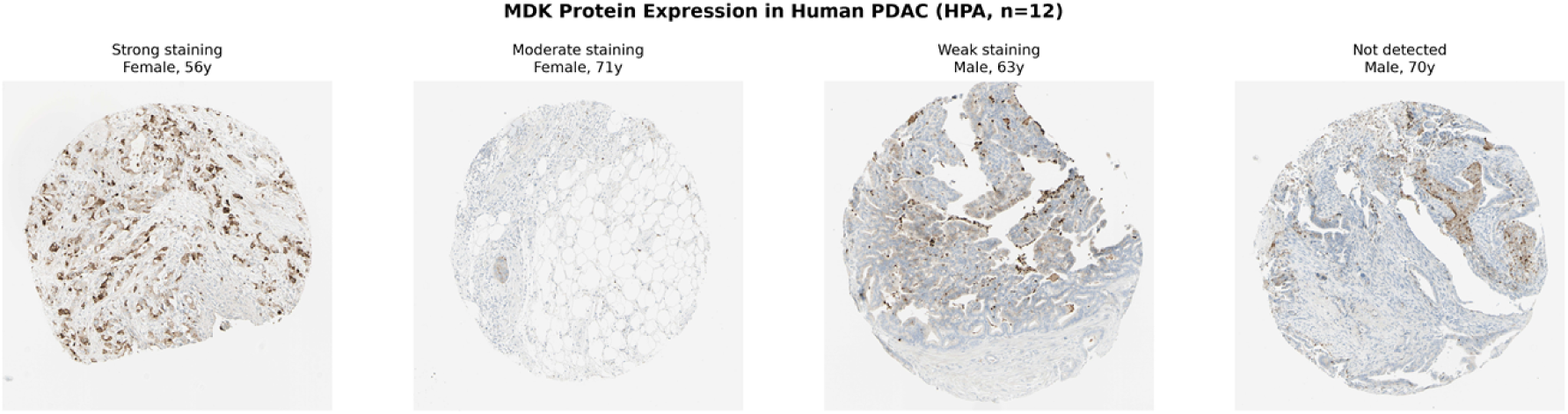
Representative MDK immunohistochemistry in human PDAC (Human Protein Atlas, antibody CAB010055).

### MDK protein in human PDAC tissues

Human Protein Atlas IHC showed cytoplasmic or membranous MDK staining in 11 of 12 PDAC cases, including diffuse stromal staining consistent with extracellular deposition (Fig 5).

## Discussion

The main finding of this study is that MDK is a causal risk factor for PDAC whose activity appears to be mediated through the stromal-immune compartment rather than through tumor cell-intrinsic mechanisms. This conclusion rests on several independent lines of evidence. Mendelian randomization identified MDK as the only gene among 34 candidates to pass correction for PDAC risk, with concordant direction across five methods and five GWAS. MDK was overexpressed in tumors and associated with shorter survival in an independent cohort. Multi-omics factor analysis placed MDK within an immune-excluded factor marked by TLS gene expression but deficient antigen-presenting cells. Single-cell data and published functional work localized MDK production to stromal cells, and CRISPR screening showed no tumor cell fitness effect.

A point that deserves emphasis is the specific character of the immune factor to which MDK belongs. This factor does not represent a generally inflamed tumor. Rather, it combines TLS gene expression with depletion of dendritic cells and macrophages, a configuration that matches the immune-excluded phenotype in which peritumoral lymphoid structures form without supporting effective intratumoral T cell infiltration [3,13]. Given the established activities of MDK as an inhibitor of dendritic cell maturation and a promoter of M2 polarization [6,18], a plausible interpretation is that stromal MDK contributes to the antigen-presentation defect that limits productive antitumor immunity. The association of MDK upregulation with SMAD4 inactivation, observed at both the transcript and the genomic level, provides a link to a core tumor suppressor pathway in PDAC.

These observations have translational implications. Current treatments for PDAC, including chemotherapy, KRAS inhibitors, and checkpoint blockade [23], largely target tumor cells or T cells without addressing the myeloid and stromal barriers that exclude immune effectors [9,10]. Preclinical studies have shown that MDK-neutralizing antibodies and MDK-targeted delivery platforms can reduce tumor growth, reprogram fibroblasts, and improve vascular perfusion [11,12,16]. Whether MDK blockade can enhance antitumor immunity in PDAC remains to be tested experimentally.

### Limitations

Several limitations should be kept in mind. The colocalization probability (PP.H4 = 0.24) is below the conventional threshold, reflecting the limited power of the FinnGen PDAC GWAS (1,252 cases); larger cohorts such as PanScan-PanC4 would be needed for definitive colocalization. Nevertheless, the primary inverse-variance-weighted estimate itself was well powered: with the observed instrument strength, the study had 80% power to detect an odds ratio of approximately 1.3, and power to detect the observed effect size (OR = 1.51) exceeded 99%. No functional experiments were performed; the proposed mechanism rests on published studies and on the internal consistency of the multi-omics, immune, single-cell, and CRISPR data. The survival association was significant in GSE62452 but not in TCGA-PAAD, and although meta-analysis indicated no heterogeneity, MDK should not yet be regarded as a validated prognostic biomarker. Finally, single-cell dropout limits the precision of cell-type assignment for a secreted factor. In addition, our findings derive mainly from populations of European ancestry and from TCGA, and their generalizability to other ancestral groups and to other tumor types in which MDK may have distinct functions remains to be established.

## Conclusions

In conclusion, our analyses identify MDK as a causal, microenvironment-mediated factor in PDAC, produced by stromal cells and associated with an immune-excluded phenotype. These results provide a rationale for testing whether MDK blockade can restore antitumor immunity in this difficult-to-treat disease.

## Supporting information

S1 Fig. MDK expression by tumor stage and histological grade in GSE62452.

S2 Fig. MDK expression by SMAD4 genomic status in TCGA-PAAD.

S1 Table. Mendelian randomization results for MDK across five methods.

## Ethics statement

Ethical approval was not required for this study, which used only publicly available, de-identified data from GTEx, FinnGen, TCGA, GEO, DepMap, and the Human Protein Atlas.

## Data availability

All data are publicly available: GTEx v8 (https://gtexportal.org/), FinnGen R13 (https://www.finngen.fi/), TCGA-PAAD via cBioPortal (https://www.cbioportal.org/), GSE62452 and GSE235449 via GEO (https://www.ncbi.nlm.nih.gov/geo/), DepMap Public 25Q3 (https://depmap.org/portal/), and the Human Protein Atlas (https://www.proteinatlas.org/). Analysis code is available from the corresponding author on request.

## Author contributions

Conceptualization: Lishan Xu, Zhengchen Yang, Xuejian Chen. Methodology: Zhengchen Yang, Xuejian Chen. Formal analysis: Zhengchen Yang. Writing – original draft: Zhengchen Yang, Xuejian Chen. Writing – review & editing: Lishan Xu, Zhengchen Yang, Xuejian Chen, Zihao Hu. Supervision: Lishan Xu. Funding acquisition: Lishan Xu.

---

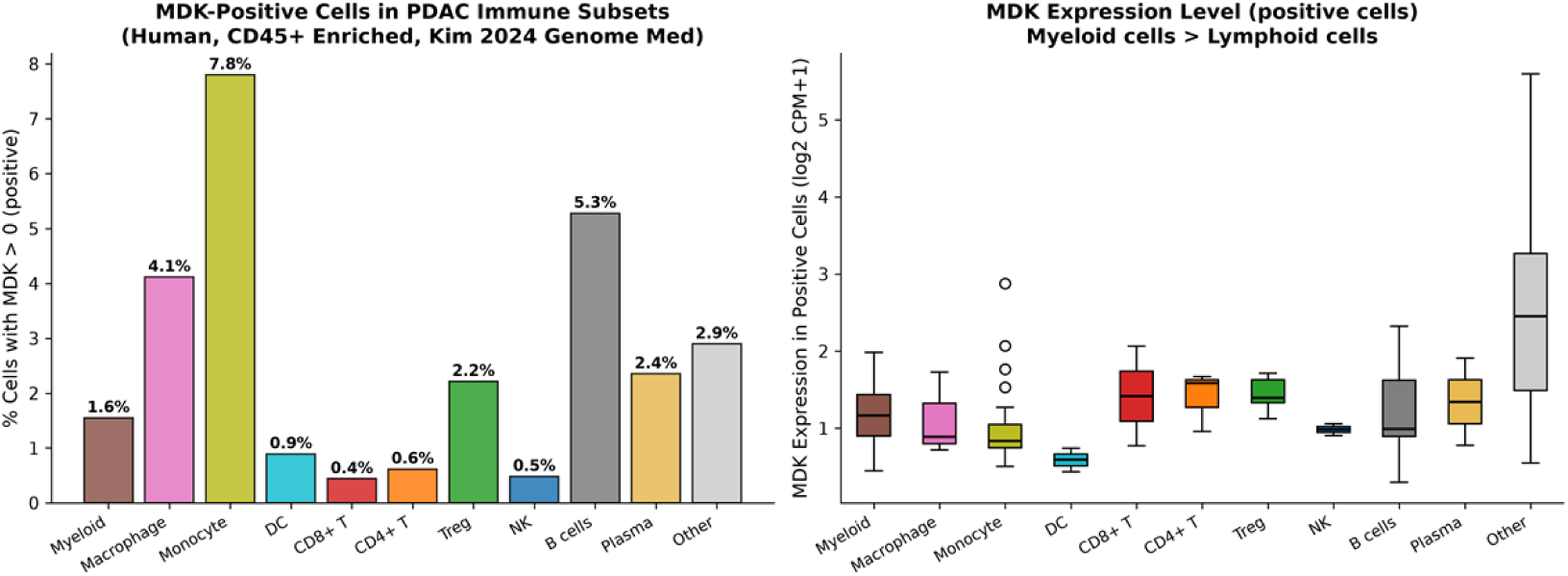

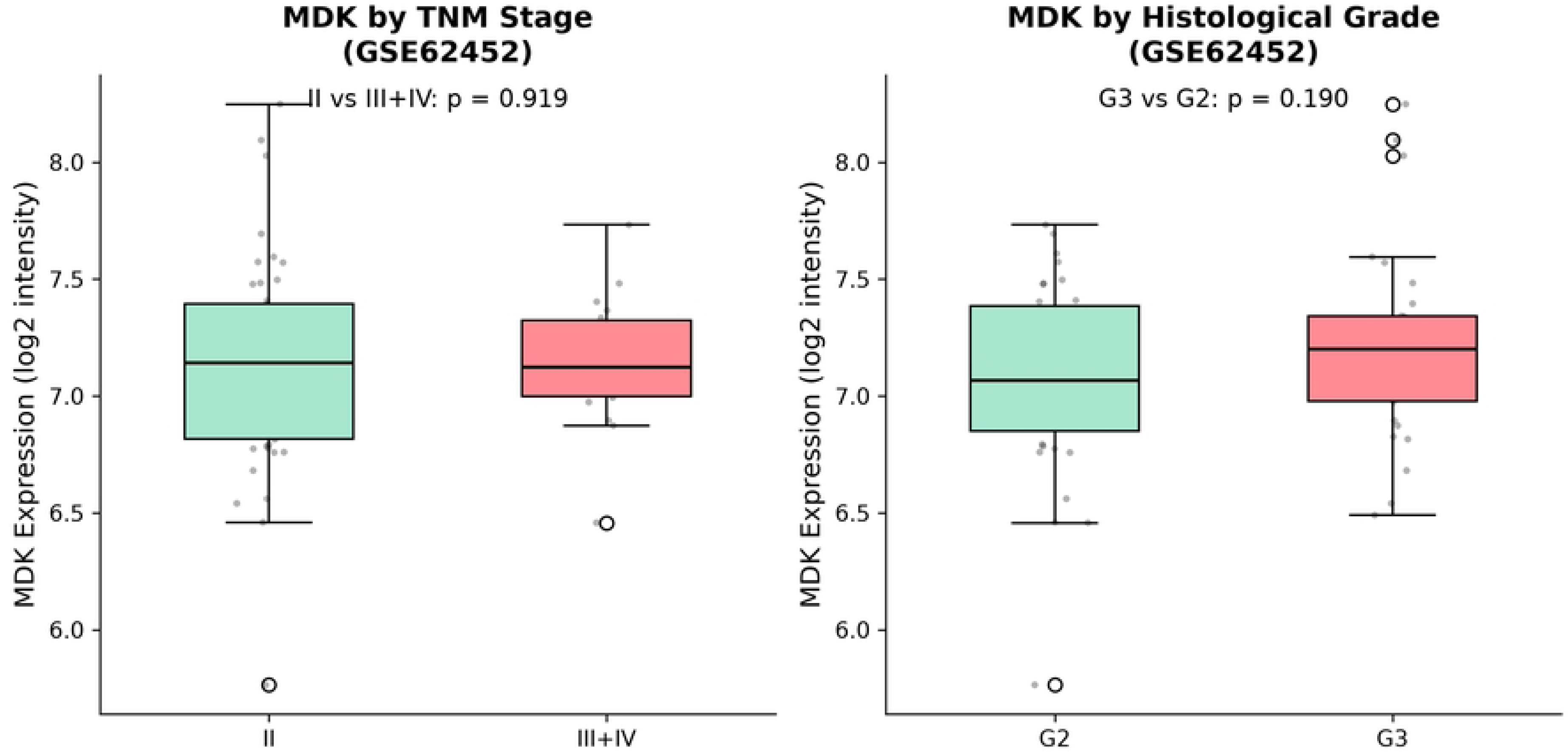

